# *Schizosaccharomyces* orthogroup (SOG) resource: a web platform for exploring gene conservation in fission yeasts

**DOI:** 10.64898/2026.01.24.701316

**Authors:** Guo-Song Jia, Fang Suo, Ambre Noly, Philippe Fort, Yue Liang, Wen Li, Wen-Cai Zhang, Heng-Le Li, Xiao-Min Du, Fan-Yi Zhang, Tong-Yang Du, Yu Hua, Feng-Yan Bai, Qi-Ming Wang, Michael Brysch-Herzberg, Dominique Helmlinger, Li-Lin Du

## Abstract

The fission yeast *Schizosaccharomyces pombe* is a prominent model organism widely used to investigate fundamental cellular mechanisms. In addition to *S. pombe*, the genus *Schizosaccharomyces* includes six other species—*S. octosporus*, *S. japonicus*, *S. cryophilus*, *S. osmophilus*, *S. lindneri*, and *S. versatilis*. These fission yeast species share a common ancestor from which the genus diversified over more than 200 million years. This extensive evolutionary divergence provides opportunities for comparative genomics. Here, we present the *Schizosaccharomyces* orthogroup (SOG) resource, a web platform developed from our high-quality genome assemblies, gene annotations, and orthology assignments. Most fission yeast genes are assigned to one of over 5,000 orthogroups. The platform enables users to visualize orthogroup sequence alignments and phylogenetic trees, retrieve coding and flanking sequences, and explore the conservation of local synteny. This resource will benefit researchers focusing on individual genes as well as those investigating gene evolution at broader scales. It is freely accessible at https://www.sogweb.org.

**TAKE AWAY:**

- The SOG resource covers all known species of *Schizosaccharomyces*.
- The platform is built on high-quality genome assemblies and annotations.
- Most genes are assigned to one of over 5,000 orthogroups.
- Users can view and explore alignments, phylogenetic trees, and local synteny.
- This free resource aids functional and evolutionary research.

## INTRODUCTION

Fission yeasts of the genus *Schizosaccharomyces* are an evolutionarily ancient clade of ascomycete fungi whose common ancestor existed over 200 million years ago (Rhind et al., 2011; Shen et al., 2020). This genus belongs to the subphylum *Taphrinomycotina* and comprises seven formally described species: *S. pombe*, *S. japonicus*, *S. octosporus*, *S. cryophilus*, *S. osmophilus*, *S. lindneri*, and *S. versatilis* (Helston, Box, Tang, & Baumann, 2010; Vaughan-Martini & Martini, 2011; Brysch-Herzberg et al., 2019; Brysch-Herzberg et al., 2023; Brysch-Herzberg et al., 2024). Among them, *S. pombe* has served as a pivotal model organism for over seven decades, driving groundbreaking discoveries across diverse fields, including cell cycle regulation, cell morphogenesis, epigenetics, and genome integrity maintenance (Hoffman, Wood, & Fantes, 2015; Hayles & Nurse, 2018). More recently, *S. japonicus* and *S. octosporus* have emerged as comparative models for evolutionary developmental cell biology and the study of killer meiotic drivers (Klar, 2013; Aoki, Furuya, & Niki, 2017; Seike, Shimoda, & Niki, 2019; Chapman, Taglini, & Bayne, 2022; De Carvalho et al., 2022; Rutherford, Harris, Oliferenko, & Wood, 2022; Alam, Gu, Reichert, Bähler, & Oliferenko, 2023; Gu, Alam, & Oliferenko, 2023; Rao et al., 2025).

Building on the extensive knowledge base of *S. pombe*, the considerable evolutionary divergence among fission yeast species offers an exceptional opportunity for comparative genomics, as demonstrated by the pioneering study of Rhind et al. that compared the genomic makeups of *S. pombe*, *S. japonicus*, *S. octosporus*, and *S. cryophilus* (Rhind et al., 2011). To further explore genomic diversity across the genus, we undertook global sampling and extensive genome sequencing of fission yeasts (Brysch-Herzberg, Jia, Seidel, Assali, & Du, 2022; Jia et al., unpublished manuscript). This effort uncovered two distinct subspecies-level lineages—*S. pombe* CN and *S. osmophilus* CA—which separated millions of years ago from the known lineages of their respective species (Jia et al., unpublished manuscript). Representative strains of the nine species/lineages of *Schizosaccharomyces* constitute a diverse genus-wide resource ideal for comparative analysis of gene and genome evolution (Figure 1A).

**Figure 1.**
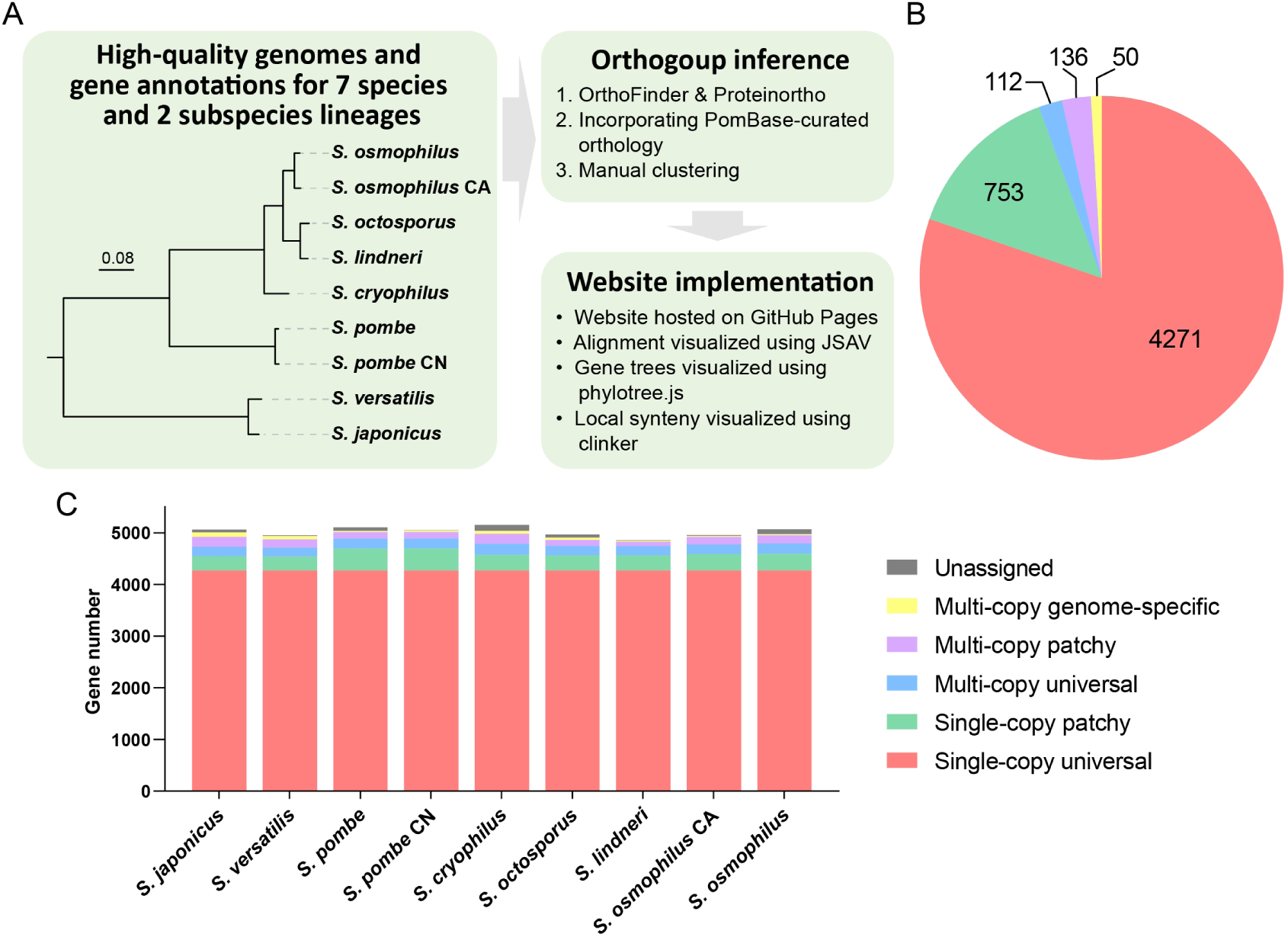
Overview of the *Schizosaccharomyces* orthogroup (SOG) resource. (A) Schematic workflow for SOG resource construction. The pipeline integrates high-quality genome assemblies and annotations from nine reference strains—representing seven *Schizosaccharomyces* species and two newly discovered lineages (*S. pombe* CN and *S. osmophilus* CA)—to define orthogroups. Initial orthogroup inference was performed using OrthoFinder and Proteinortho, followed by refinement based on PomBase-curated *S. japonicus*–*S. pombe* orthology relationships and additional manual corrections. The final SOG set (5,322 orthogroups) powers a lightweight, client-side web platform built with HTML and JavaScript and deployed via GitHub Pages, enabling interactive visualization of multiple sequence alignments (via JSAV), phylogenetic trees (via phylotree.js), and local synteny (via clinker). (B) Distribution of SOG types. The 5,322 orthogroups are classified into five categories: 4,271 single-copy universal (red), 753 single-copy patchy (green), 112 multi-copy universal (blue), 136 multi-copy patchy (purple), and 50 multi-copy genome-restricted (yellow). (C) Gene content per genome by SOG type. Each stacked bar represents one of the nine genomes; segment colors correspond to the SOG types in (B), with unassigned genes shown in grey.

To maximize the value of this resource, we generated high-quality chromosome-level genome assemblies for representative strains of all nine species/lineages and performed comprehensive gene annotations (Jia et al., 2023; Brysch-Herzberg et al., 2024; Li et al., unpublished manuscript; Jia et al., unpublished manuscript). Leveraging these data, we inferred orthologous relationships of protein-coding genes across the genus, and, as a service to the fission yeast community, developed the *Schizosaccharomyces* orthogroup (SOG) resource. This resource is a web-accessible platform that integrates multiple modules: alignments of amino acid sequences, gene trees, local synteny plots, and downloadable nucleotide sequences of coding DNA sequences (CDSs) and CDS-flanking regions. It enables researchers to examine cross-species evolutionary conservation and divergence, supporting both targeted gene studies and broad systematic inquiry.

## MATERIALS AND METHODS

### Genome assemblies and protein-coding gene annotation

The SOG resource is based on nine chromosome-level genome assemblies representing all seven recognized fission yeast species and two newly identified divergent lineages. The assemblies for *S. osmophilus*, *S. japonicus*, and *S. versatilis* are those recently published by us (Jia et al., 2023; Brysch-Herzberg et al., 2024). The other six genome assemblies are newly generated and will be detailed in upcoming publications (Li et al., unpublished manuscript; Jia et al., unpublished manuscript). All assemblies consist of fully contiguous chromosomes that terminate either in telomeric repeats or rDNA arrays.

Protein-coding gene annotation was performed for all nine genome assemblies. Existing gene annotations for *S. pombe*, *S. octosporus*, *S. cryophilus*, *S. japonicus*, and *S. osmophilus* were either retained or transferred to the new assemblies (Rhind et al., 2011; Rutherford et al., 2022; Jia et al., 2023; Rutherford, Lera-Ramírez, & Wood, 2024). For *S. versatilis*, *S. lindneri*, *S. osmophilus* CA, and *S. pombe* CN, annotations were transferred from the closest relatives (*S. japonicus* to *S. versatilis*; *S. octosporus* to *S. lindneri*; *S. osmophilus* to *S. osmophilus* CA; *S. pombe* to *S. pombe* CN). Gene prediction was conducted to complement the annotation transfer. Rigorous manual curation was performed to validate the annotations and correct errors. Full details of the annotation procedures and results will be described in forthcoming publications (Li et al., unpublished manuscript; Jia et al., unpublished manuscript).

To facilitate cross-referencing, we preserved existing systematic IDs whenever possible. For *S. octosporus*, *S. cryophilus*, and *S. japonicus*, where a locus_tag prefix is part of the commonly used systematic IDs, we changed the locus_tag prefix while retaining the unique serial number that follows the prefix and underscore (e.g., from SJAG_03048 to SZJAPO_03048).

### Orthogroup inference

Orthogroups across nine genomes were inferred using two complementary tools: OrthoFinder (v2.5.4) and Proteinortho (v6.1.0) (Lechner et al., 2011; Emms & Kelly, 2019). An important distinction between these two tools is that only Proteinortho utilizes synteny information to infer orthology, which is implemented through its PoFF extension (Lechner et al., 2014).

Our evaluations indicated that these two tools have complementary strengths: OrthoFinder excels at delineating large gene families (containing dozens of members), while Proteinortho tends to generate smaller groups. Crucially, Proteinortho leverages synteny to detect orthology that is not apparent based solely on sequence similarity. This capability is exemplified by its ability to resolve the orthologous relationships of ribosomal protein genes, which duplicated in the fission yeast common ancestor but underwent sequence homogenization after lineage splits (Mullis et al., 2020).

To capitalize on these strengths, we constructed an orthogroup set primarily from OrthoFinder output, with targeted refinement using Proteinortho results. Specifically, we re-examined OrthoFinder-generated multi-copy orthogroups that contained the same number of genes in each genome. In instances where Proteinortho resolved such groups into single-copy orthogroups, the original OrthoFinder groups were split accordingly. This strategy yielded 142 additional single-copy universal orthogroups: 124 derived from groups with two genes per genome, 9 from groups with three, and 9 from larger groups (one with four and one with five genes per genome). Manual inspection confirmed that these splits were supported by conserved local synteny.

The high evolutionary divergence of *S. japonicus* and *S. versatilis* from other fission yeast species poses challenges for orthology detection. To address this, we augmented orthology assignment using PomBase-curated *S. japonicus*–*S. pombe* orthologs (https://www.pombase.org/monthly_releases/2025/pombase-2025-12-01/curated_orthologs/, accessed: 2025-09-26) (Rutherford et al., 2022). A total of 109 PomBase-curated one-to-one *S. japonicus*–*S. pombe* ortholog pairs were not detected in our initial orthogroup assignments. We verified 102 pairs based on manual inspection of local synteny and phylogenetic relationships and used them to merge orthogroups.

Genes not assigned to any orthogroups were individually assessed manually, resulting in the addition of 12 orthogroups that were missed in the previous workflow. The finalized SOGs (version 1.0) are shown in Supplementary Table S1.

### Chromosomal map visualization

To visualize the chromosomal locations of gene members within each SOG, we generated chromosomal maps using NGenomeSyn (He et al., 2023). The chromosomes of the nine genomes are displayed as horizontal grey bars arranged along nine parallel tracks. Global synteny was visualized by drawing light grey lines between orthologous gene pairs (single-copy universal SOGs) on adjacent tracks. Centromeres and 18S–5.8S–28S rDNA arrays were annotated on the chromosome bars. The genomic positions of gene members within a given SOG were projected onto this layout and indicated by red vertical lines. Finally, the exported SVG files were adjusted to resolve text overlaps and enhance legibility.

### Multiple sequence alignment and phylogenetic analysis

Amino acid sequences were aligned using MAFFT (v7.475) under the E-INS-i mode (Katoh & Standley, 2013). Phylogenetic trees were then inferred from these alignments using IQ-TREE (v2.0.3) with the following options: -m TEST -bb 1000 -alrt 1000 -nt 1 (Minh et al., 2020). All phylogenetic trees were midpoint-rooted using the ape package in R (Paradis & Schliep, 2019). Alignment visualization on the web platform was implemented using the JSAV library (Karavirta & Shaffer, 2013). Phylogenetic tree visualization was implemented using phylotree.js (Shank, Weaver, & Kosakovsky Pond, 2018).

### Integrating PomBase-curated *Saccharomyces cerevisiae* and human orthologs into protein sequence alignments

PomBase-curated *S. pombe*–*Saccharomyces cerevisiae* orthologs and *S. pombe*–*Homo sapiens* orthologs were obtained from PomBase (https://www.pombase.org/monthly_releases/2025/pombase-2025-12-01/curated_orthologs/, accessed: 2025-12-08).

The *S. cerevisiae* proteome, containing 6,722 sequences, was obtained from the *Saccharomyces* Genome Database (SGD; http://sgd-archive.yeastgenome.org/sequence/S288C_reference/orf_protein/orf_trans_all.fasta.gz accessed: 2025-12-10). Because the canonical S288C strain lacks several α-galactosidase paralogs, we supplemented this set with four additional proteins: Mel1 (UniProt: P04824), Mel2 (UniProt: P41945), Mel5 (UniProt: P41946), and Mel6 (UniProt: P41947). These four sequences correspond to loci YSC0019, YSC0021, YSC0024, and YSC0025 in non-reference strains and were retrieved individually from UniProt. For *H. sapiens,* the reference proteome consisting of manually reviewed (Swiss-Prot) canonical entries was obtained from UniProt Proteomes (Proteome ID UP000005640; https://www.uniprot.org/proteomes/UP000005640; accessed: 2025-12-10). This dataset comprised 20,405 sequences.

To ensure alignment reliability, we filtered the orthologs using reciprocal BLASTP (NCBI BLAST+ v2.5.0, default parameters). Each *S. pombe* protein was queried against the full proteome of the target species and vice versa. Only ortholog pairs that satisfied the reciprocal hit criteria (E-values no larger than 1E-2 in both directions) were retained. After filtering, 4,828 *S. pombe*–*Saccharomyces cerevisiae* protein pairs (Supplementary Table S2) and 4,809 *S. pombe*–*Homo sapiens* protein pairs remained (Supplementary Table S3). The *S. cerevisiae* and *H. sapiens* protein sequences were added to the existing SOG alignments using the --add option in MAFFT (v7.475).

### Local synteny visualization for single-copy SOGs

Local synteny conservation across all single-copy orthogroups was visualized using clinker (v0.0.31) (Gilchrist & Chooi, 2021). For each SOG, a set of GenBank (.gbk) format files was generated; each file contained the DNA sequence and CDS annotations for the genomic region centered on the respective SOG member, spanning five upstream and five downstream flanking genes. These files were provided to clinker, which performed all-against-all pairwise global alignments of amino acid sequences (using Biopython’s Bio.Align.PairwiseAligner) and calculated sequence identities.

We evaluated a series of identity thresholds (0.18 to 0.30, in 0.01 increments) as the minimum cutoff for grouping genes into similarity groups (Figure S1). Only genes belonging to the same similarity group are connected by links in the visual display. We chose the threshold of 0.25 to balance two competing objectives: (1) maximizing the number of links between orthologs and (2) minimizing spurious links between non-homologous genes.

The HTML outputs of clinker were adjusted to minimize file size and optimize web performance.

### Web platform implementation

The SOG resource was developed as a static web application to ensure long-term accessibility and minimal maintenance requirements. The user interface was built using standard HTML, CSS, and JavaScript. The platform is deployed statically via GitHub Pages and requires no dedicated backend server. This architecture ensures high availability through GitHub’s content delivery network. The source code, including the HTML and JavaScript files used to generate the static site, is available on GitHub at https://github.com/fsnibs10/SOG. Users can access the SOG resource at https://www.sogweb.org.

### Disclosure on the use of artificial intelligence tools

The manuscript was proofread using the artificial intelligence tools editGPT, Qwen3-Max, and Gemini 3 Pro Preview. After using these tools, the authors reviewed and edited the content as needed and take full responsibility for the content of the published article.

## RESULTS AND DISCUSSION

### Assigning fission yeast protein-coding genes to SOGs

To enhance understanding of gene conservation and divergence across the *Schizosaccharomyces* genus, we assigned fission yeast protein-coding genes to *Schizosaccharomyces* orthogroups (SOGs), where each SOG comprises orthologous genes descended from a single ancestral gene. To ensure high-quality orthogroup inference, we employed a workflow integrating state-of-the-art computational methods, PomBase-curated *S. japonicus*–*S. pombe* orthology relationships, and manual validation (Figure 1A).

We inferred a total of 5,322 SOGs. Based on the number and distribution of SOG members across genomes, these orthogroups were classified into five categories (Figure 1B): 4,271 single-copy universal SOGs (one gene in each genome); 753 single-copy patchy SOGs (one gene in some genomes and absent in others); 112 multi-copy universal SOGs (present in all genomes, with paralogs in at least one); 136 multi-copy patchy SOGs (present in some genomes, with paralogs in at least one); and 50 multi-copy genome-specific SOGs (present in only one genome, with paralogs) (Figure 1B).

In each of the nine genomes, more than 80% of protein-coding genes belong to single-copy universal SOGs (Figure 1C and Table S4), reflecting strong conservation of both gene content and copy number over more than 200 million years of *Schizosaccharomyces* evolution. Among the remaining genes, a significant proportion (35–54%) belongs to single-copy patchy SOGs, a pattern attributable to lineage-specific gene loss or gain.

Notably, *S. pombe* and *S. pombe* CN harbor a disproportionately high number of genes in single-copy patchy SOGs compared to the other genomes. This excess is largely driven by genes present exclusively in this pair. This pattern may partly stem from differences in the criteria used to filter low-confidence gene models: 17% of the genes exclusive to this pair are labeled as “dubious” in PomBase (Rutherford et al., 2024), suggesting that inclusion of gene models with limited supporting evidence contributes to the inflated count.

Genes belonging to multi-copy SOGs account for 6–9% of the genes in individual genomes. The SOG with the highest member count is SOG_5586 (90 members), which corresponds to retrotransposons belonging to the Tf1/sushi group (Butler, Goodwin, Simpson, Singh, & Poulter, 2001; Goodwin & Poulter, 2001; Jia et al., 2023). The ability of retrotransposons to mobilize via a “copy-and-paste” mechanism likely underlies their high copy number. Ranked second and third are SOG_5587 (*hsp16* genes encoding small heat shock proteins, 70 members) and SOG_5588 (encoding hexose transporters, 55 members). The elevated copy number of hexose transporter genes has been proposed to reflect adaptation to glucose-rich environments (Lin & Li, 2011). The exceptionally high copy number of *hsp16* genes is an intriguing phenomenon and will be described in more detail in a later section.

Genes not assigned to any SOG (“unassigned” in Figure 1C and Table S4) constitute no more than 2% of the genes in any individual genome. These are likely enriched for low-confidence gene models. Genomes with pre-existing annotations (*S. pombe*, *S. japonicus*, *S. cryophilus*, *S. octosporus*, and *S. osmophilus*) contained more unassigned genes than the four newly annotated genomes (*S. versatilis*, *S. lindneri*, *S. osmophilus* CA, and *S. pombe* CN), likely due to the stringent filtering criteria applied during our annotation of the latter four.

### Architecture and implementation of the SOG web platform

To provide open and user-friendly access to SOG-based datasets, we developed the SOG web resource as a lightweight, client-side application. Built with HTML and JavaScript, the platform is deployed statically via GitHub Pages and requires no dedicated backend server (Figure 1A and Figure 2). The platform’s entry point is a searchable table listing all SOGs and their affiliated genes, where each SOG ID links to a dedicated SOG page. These SOG pages contain download links for member coding sequences (with and without introns) and sequences of 1-kb upstream/downstream flanking regions, alongside a plot showing the chromosomal positions of SOG members across the nine genomes.

**Figure 2.**
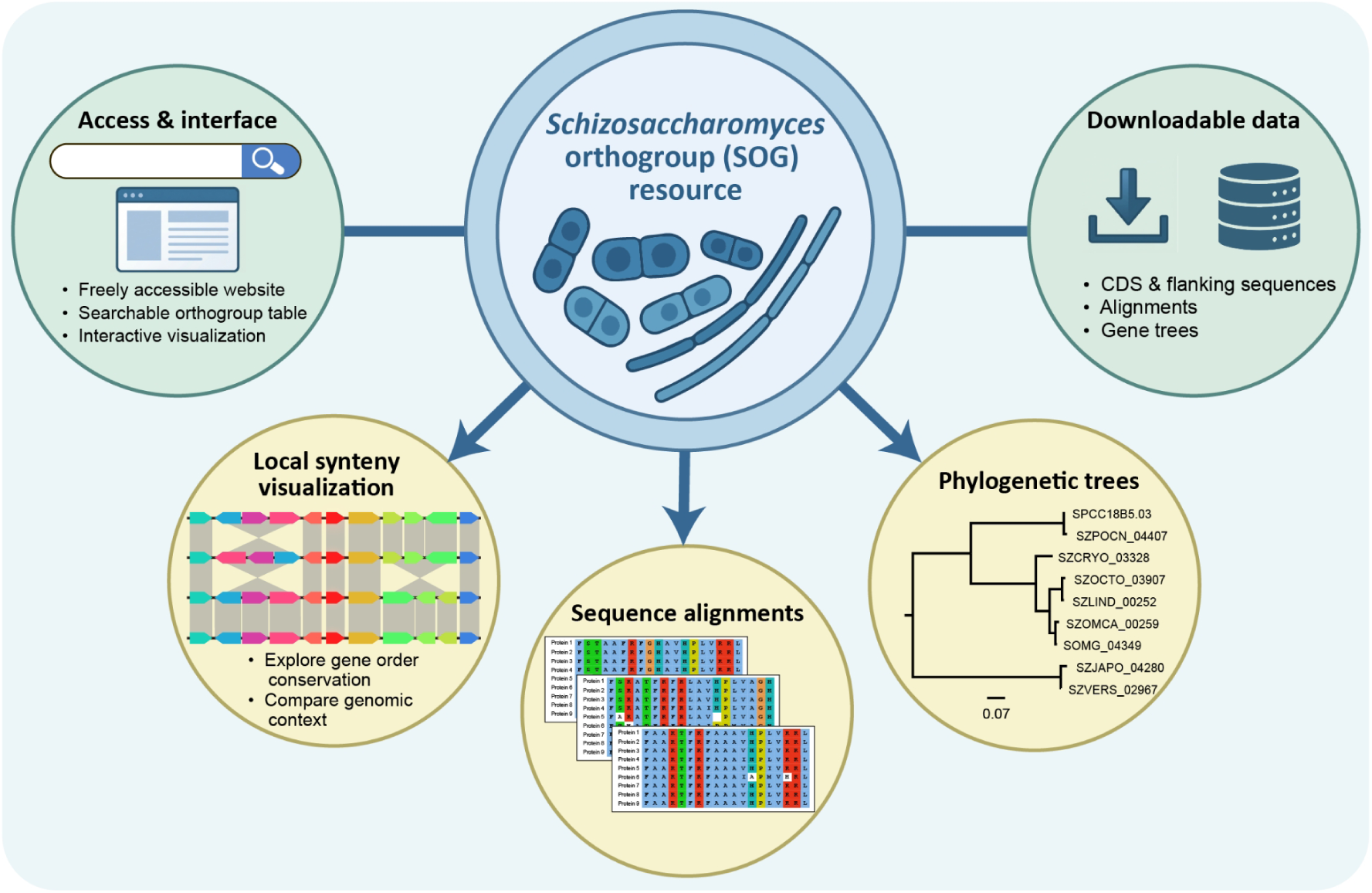
Architecture of the SOG web platform. The searchable SOG table functions as the navigation hub, linking to downloadable datasets and three interactive visualization modules: (1) sequence alignments, (2) phylogenetic trees, and (3) local synteny. The platform is implemented entirely in static HTML/JavaScript and requires no backend server.

From these pages, users can access interactive visualizations of three precomputed datasets: multiple sequence alignments (MSAs), midpoint-rooted maximum-likelihood phylogenetic trees, and local synteny plots (Figure 2). MSAs are rendered using the JSAV library (Karavirta & Shaffer, 2013). Phylogenetic trees are visualized via phylotree.js, which supports layout switching (rectangular or radial), spacing adjustment, reordering of deepest clades, and toggleable tip label alignment (Shank, Weaver, & Kosakovsky Pond, 2018). Finally, single-copy SOG synteny is displayed using clinker (Gilchrist & Chooi, 2021), which enables track reordering, sequence flipping and movement, zooming, and fine-grained control over visualization parameters.

Together, these features provide an integrated, interactive environment for exploring orthogroup evolution across the *Schizosaccharomyces* genus. We illustrate the utility of the SOG resource through two use cases below.

### Use case 1: exploring the evolution of *cdc2*

Single-copy SOGs—including both single-copy universal and single-copy patchy SOGs—account for 90–94% of protein-coding genes in individual genomes. To demonstrate how the SOG platform facilitates the evolutionary exploration of these genes, we present a use case regarding the *S. pombe* gene *cdc2*—a historically significant and extensively characterized gene (Figures 3 and 4).

**Figure 3.**
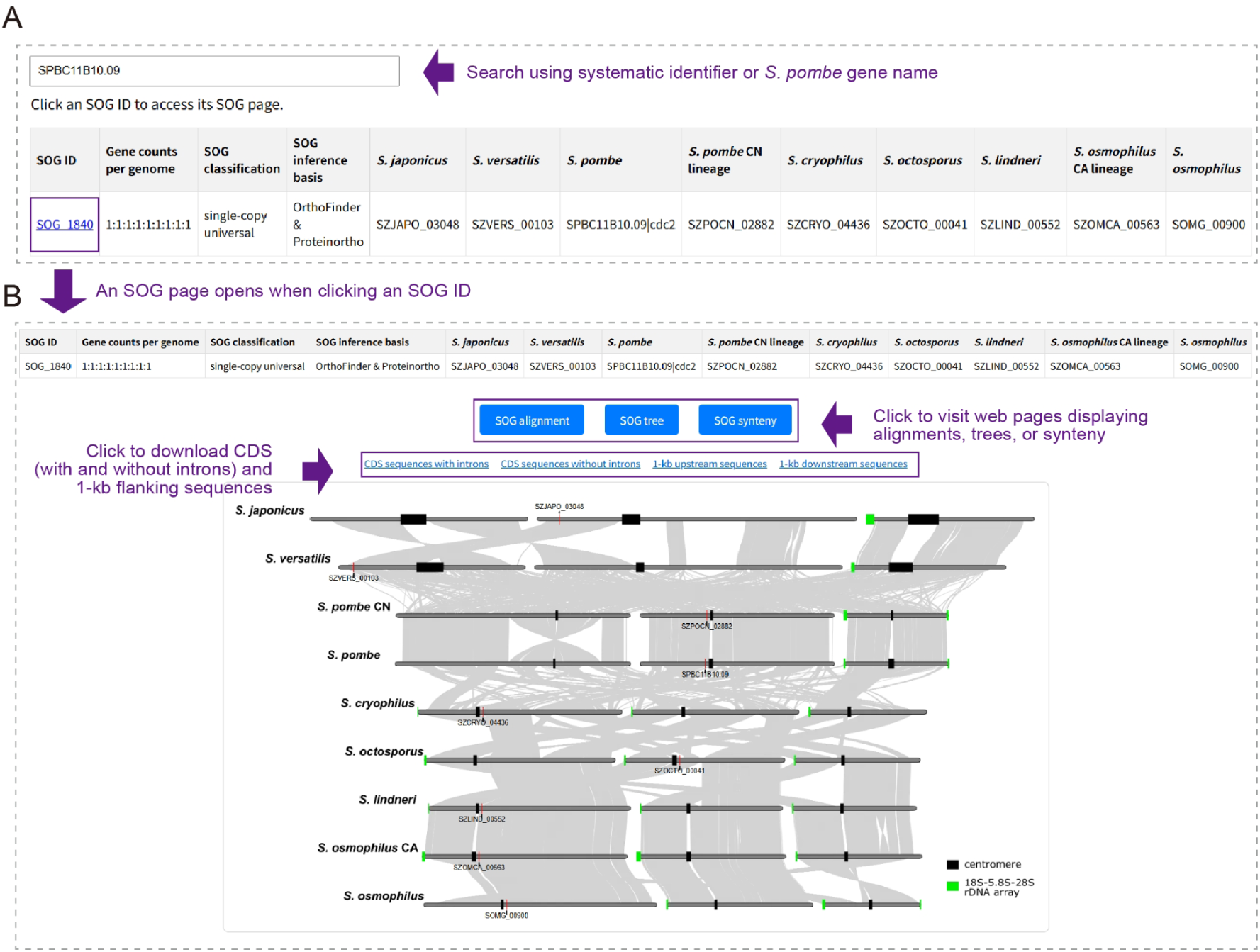
Querying a gene of interest in the SOG database (example: *cdc2*). (A) Gene search interface. Users can input a systematic ID (e.g., “SPBC11B10.09”) or *S. pombe* gene name (e.g., “cdc2”) to retrieve the corresponding SOG (SOG_1840) and its orthologs across all nine genomes. (B) The SOG page of SOG_1840. This interface provides download links for CDSs (with and without introns) and 1-kb upstream/downstream flanking sequences, as well as buttons to launch the “SOG alignment,” “SOG tree,” and “SOG synteny” modules. A genome-wide synteny plot displays the chromosomal positions of orthologous genes as red vertical lines. Grey bars represent chromosomes, while black and green blocks denote centromeres and rDNA arrays, respectively.

**Figure 4.**
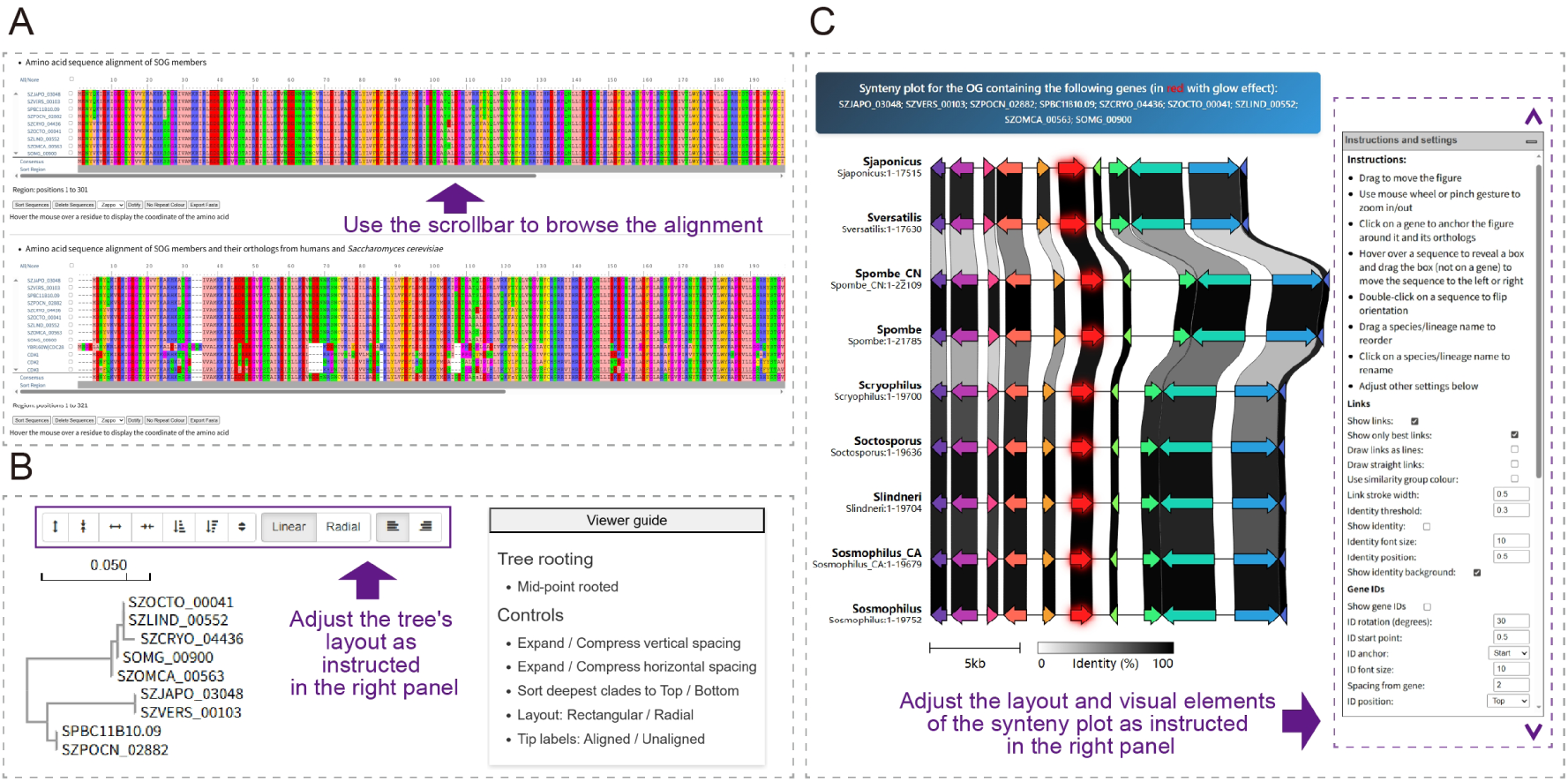
Visualization modules of the SOG platform (example: SOG_1840). (A) Sequence alignment viewer. Top: Multiple sequence alignment of SOG members. Bottom: When orthologs from *Saccharomyces cerevisiae* or *Homo sapiens* are available, a separate alignment including these sequences is displayed. (B) Phylogenetic tree viewer. A midpoint-rooted maximum-likelihood tree is rendered with interactive controls. A side panel explains customization options. (C) Local synteny viewer. For single-copy SOGs, genomic tracks are displayed covering an 11-gene window (five upstream, the SOG member, and five downstream). Genes are represented as arrows. The SOG member is highlighted in red with a glow effect, while flanking genes are colored by similarity group (≥25% amino acid identity). Inter-track links connect genes belonging to the same similarity group. A panel on the right explains how to adjust visualization and provides fine-grained control of visual elements.

On the SOG table page, a user can enter "cdc2" or its systematic ID (SPBC11B10.09) into the search box to locate its corresponding orthogroup, SOG_1840 (Figure 3A). Clicking the SOG ID opens the dedicated SOG page (Figure 3B), from which the following analyses can be performed:

1. Genomic location inspection: A genome-level synteny plot displays the chromosomal positions of *cdc2* orthologs across the nine *Schizosaccharomyces* genomes. Red vertical lines indicate these positions, revealing that *cdc2* resides near a centromere in most species, with the exception of *S. japonicus* and *S. versatilis*.
2. Sequence retrieval: Users can download nucleotide sequences of the coding sequences (CDSs) for all nine *cdc2* orthologs, as well as their 1- kb upstream and downstream flanking regions, facilitating analyses of regulatory cis-elements.
3. Protein sequence conservation: Clicking the “SOG alignment” button opens an interactive multiple sequence alignment (MSA) page (Figure 4A). Two alignments are provided: one containing only the *Schizosaccharomyces* SOG members, and a second that includes orthologs from *Saccharomyces cerevisiae* and *Homo sapiens*. Visual inspection of *cdc2* alignments reveals high overall sequence conservation, consistent with strong purifying selection. As expected, sequence divergence and gap frequency increase with evolutionary distance, highlighting regions of the protein that tolerate changes.
4. Phylogenetic analysis: The “SOG tree” button leads to a midpoint-rooted maximum-likelihood gene tree revealing the phylogenetic relationships among SOG members (Figure 4B). For *cdc2*, the tree exhibits a topology that largely recapitulates the species phylogeny but shows minor discordances, likely due to limited phylogenetic signal in such highly conserved sequences.
5. Microsynteny assessment: The “SOG synteny” button opens a local synteny view (Figure 4C), displaying nine tracks centered on each *cdc2* ortholog. Each track spans 11 consecutive genes (five upstream, the SOG member, and five downstream), represented as arrows. The SOG members are highlighted in red with a glow effect; flanking genes are colored by similarity group (defined at ≥25% amino acid identity), with inter-track links connecting genes belonging to the same similarity group. In the case of *cdc2*, the surrounding genes exhibit perfectly conserved order and orientation across all nine genomes, indicating strong microsynteny conservation.

This use case illustrates how the SOG platform streamlines orthogroup-based analysis by integrating alignments, trees, synteny plots, and downloadable sequences into a unified, interactive interface. It enables researchers to rapidly assess the evolutionary trajectory of specific genes. In the case of *cdc2* orthologs, visual inspection of sequence alignments and local gene order provides strong support for orthology and reveals extensive conservation consistent with strong purifying selection across the *Schizosaccharomyces* genus.

### Use case 2: exploring the evolution of *hsp16* genes

As noted earlier, *hsp16* genes constitute the second largest SOG (SOG_5587), with 70 members. We use these *hsp16* genes to illustrate how the SOG platform can be leveraged to investigate the evolutionary dynamics of a multi-copy orthogroup. *hsp16* genes are present in all nine fission yeast genomes (Figure 5A). However, copy number varies dramatically across species: *S. pombe* and *S. pombe* CN each possess a single copy; *S. japonicus* and *S. versatilis* each have two; and the remaining five genomes show substantial expansion—24, 8, 8, 10, and 14 copies in *S. cryophilus*, *S. octosporus*, *S. lindneri*, *S. osmophilus* CA, and *S. osmophilus*, respectively. The sole *hsp16* gene in *S. pombe* encodes a small heat shock protein (sHSP) that functions as a molecular chaperone, preventing aggregation of denatured proteins under thermal stress and contributing to thermotolerance (Taricani, Feilotter, Weaver, & Young, 2001; Hirose et al., 2005; Yoshida & Tani, 2005; Glatz et al., 2016).

**Figure 5.**
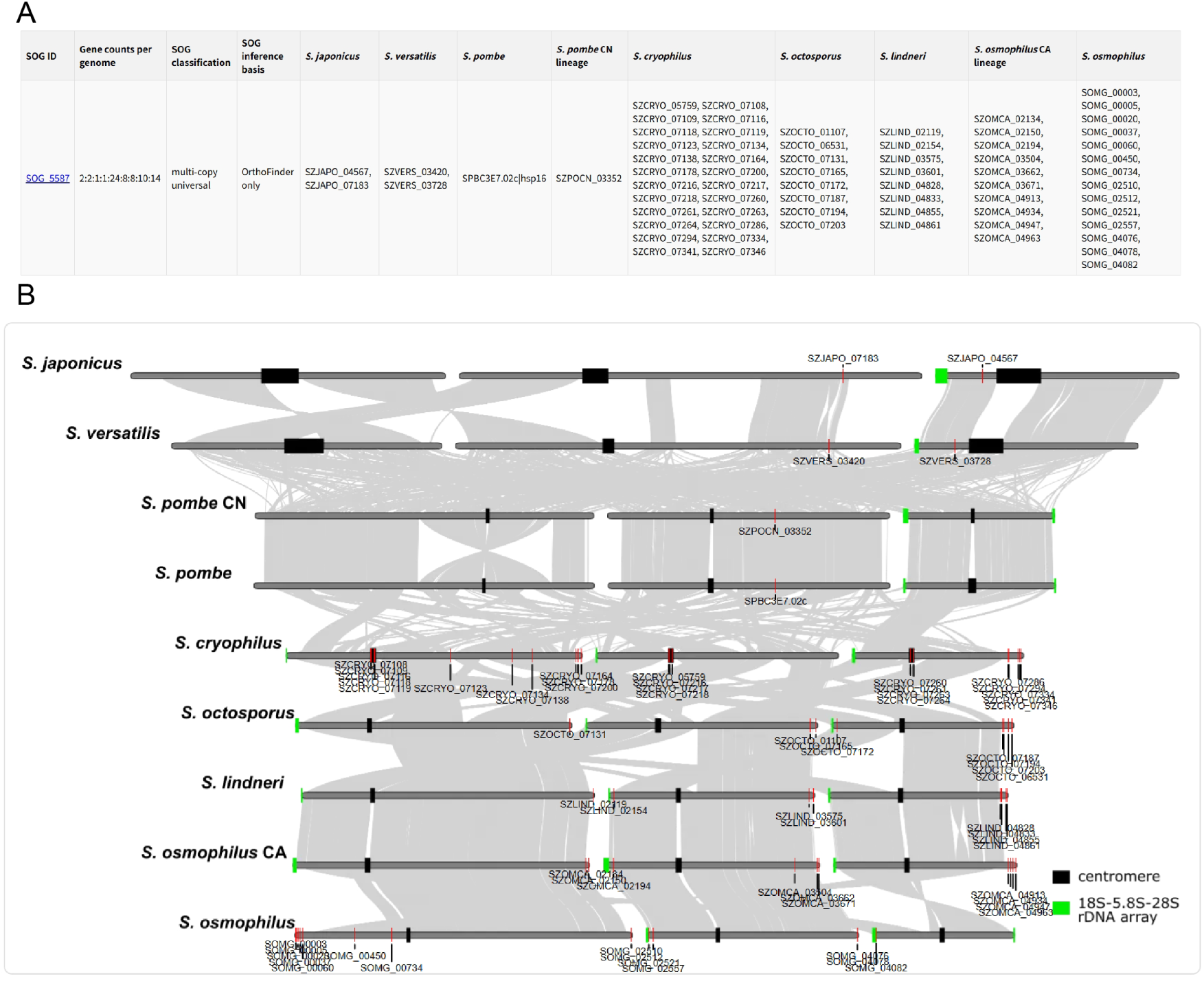
Dramatic expansion of the copy numbers of *hsp16* genes in certain fission yeast lineages. (A) SOG table entry for SOG_5587, the *hsp16* orthogroup. (B) Chromosomal locations of *hsp16* genes in the nine genomes. In genomes harboring 1–2 copies (*S. pombe*, *S. pombe* CN, *S. japonicus*, and *S. versatilis*), *hsp16* genes reside in the interior of chromosome arms. In genomes with expanded copy numbers, copies are enriched in subtelomeric regions, with *S. cryophilus* uniquely harboring centromeric copies.

The SOG platform’s genomic location plot reveals a striking correlation between copy number and chromosomal distribution (Figure 5B). In the four genomes with low copy numbers (1–2 genes), *hsp16* genes reside in the interior of chromosome arms (Figure 5B). In contrast, in the five lineages with expanded repertoires, many *hsp16* genes are concentrated in subtelomeric regions. *S. cryophilus* is unique among these; as previously reported, some of its *hsp16* genes are located within the centromeres (Tong et al., 2019).

The phylogeny of SOG_5587, accessible via the SOG tree module, displays a complex topology (Figure 6). Genes from *S. japonicus*, *S. versatilis*, *S. pombe*, *S. pombe* CN, *S. octosporus*, and *S. lindneri* largely follow patterns expected under vertical inheritance coupled with lineage-specific duplications. A duplication likely occurred in the common ancestor of *S. japonicus* and *S. versatilis*, and the two resulting paralogs diverged substantially prior to speciation—such that the two copies within each species are more dissimilar to each other than to their respective counterparts in the sister species. The *hsp16* genes of *S. octosporus* form a monophyletic clade and the *hsp16* genes of *S. lindneri* form another, consistent with independent copy number expansion after these two sibling species split.

**Figure 6.**
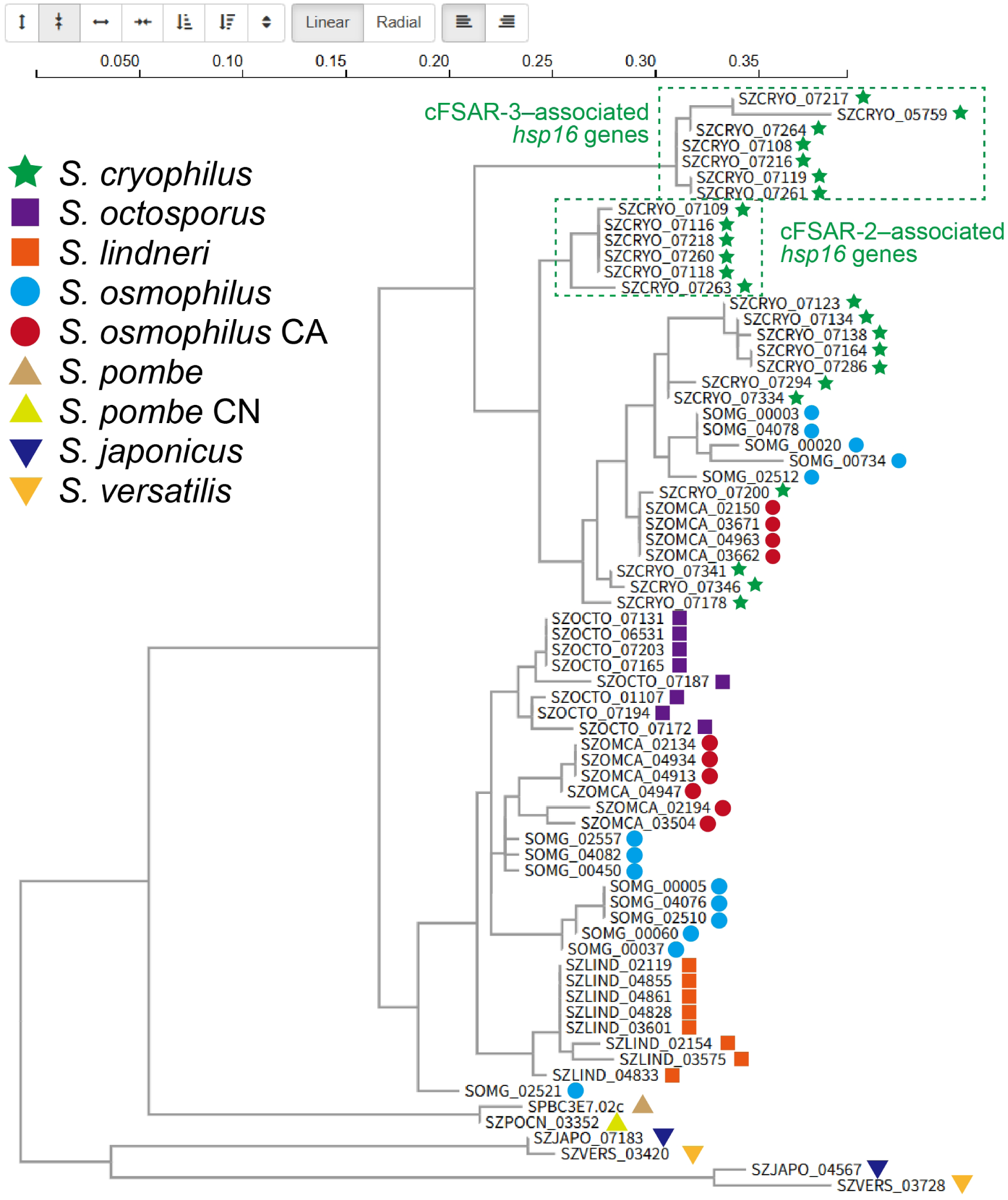
Phylogeny of *hsp16* genes reveals complex evolutionary dynamics. Maximum-likelihood phylogenetic tree of all 70 *hsp16* genes. Centromeric *hsp16* genes in *S. cryophilus* form two distinct clades corresponding to their association with cFSAR-2 or cFSAR-3 repeats, suggesting repeat-mediated paralog homogenization. Cross-species clustering—exemplified by *S. cryophilus* SZCRYO_07200 grouping with four *S. osmophilus* CA genes (SZOMCA_02150, SZOMCA_03671, SZOMCA_04963, and SZOMCA_03662)—suggests horizontal gene transfer.

However, the tree also reveals anomalies suggestive of reticulate evolution. The *hsp16* genes of *S. cryophilus* do not form a distinct clade but are intermixed with subsets of genes from *S. osmophilus* and *S. osmophilus* CA in the tree, while the rest of the latter two species’ genes cluster with those of *S. octosporus* and *S. lindneri*. These patterns imply that horizontal gene transfer (HGT) or introgression may have occurred across species boundaries.

In *S. cryophilus*, the centromere-located *hsp16* genes are embedded within two types of 5S rDNA-containing repeats, cFSAR-2 and cFSAR-3 (Tong et al., 2019). The SOG gene tree recapitulates the clustering reported by Tong et al.: cFSAR-2–associated and cFSAR-3–associated *hsp16* genes form distinct clades (Figure 6), indicating repeat-mediated homogenization.

Together, the dramatic copy number expansion, the concentration of *hsp16* genes in dynamic genomic regions (subtelomeres and centromeres), and evidence of cross-species exchange point to the action of unusual evolutionary forces on *hsp16* in certain *Schizosaccharomyces* lineages. While the nature of these forces remains unclear, we propose two non-exclusive hypotheses. First, copy number expansion may reflect adaptive responses to environmental stress. Elevated sHSP copy number has been linked to terrestrial adaptation in plants (Haslbeck & Vierling, 2015; Waters & Vierling, 2020), suggesting a potential role in niche specialization. Second, the genomic and evolutionary signatures—centromeric and subtelomeric localization, rapid diversification, and HGT—resemble those of selfish killer meiotic drivers in fission yeasts (De Carvalho et al., 2022; Xu et al., 2024; Zhang et al., 2025). This raises the possibility that *hsp16* genes have either acquired selfish properties or become co-opted into genetic conflicts involving selfish elements. Testing these hypotheses will require further experimental studies.

## Conclusion

The SOG resource establishes a comprehensive, interactive framework for exploring gene evolution across the *Schizosaccharomyces* genus. By analyzing nine high-quality genome assemblies, we classified over 5,000 orthogroups, revealing that the *Schizosaccharomyces* clade is characterized by a highly conserved core proteome: in every species, more than 80% of protein-coding genes belong to single-copy universal orthogroups. Conversely, our classification identifies specific subsets of genes subject to lineage-specific loss, gain, or expansion, offering targets for studying evolutionary innovation and adaptation.

The SOG web platform translates these genomic data into an accessible, researcher-centric tool by integrating orthogroup assignments with visualizations of sequence alignments, phylogenies, synteny, and chromosomal maps. As demonstrated by the contrasting evolutionary trajectories of the highly conserved *cdc2* and the dynamically evolving *hsp16* family, the platform supports both robust orthology validation in stable gene families and hypothesis generation for complex evolutionary scenarios involving copy number variation, horizontal transfer, and paralog homogenization. Furthermore, the platform’s client-side architecture ensures long-term accessibility without reliance on dedicated backend infrastructure, making it a sustainable community asset. Freely available at https://www.sogweb.org and soon to be linked from the “External references” section of PomBase gene pages, the SOG resource will empower diverse research directions—from functional studies of individual genes to broad investigations of genome evolution, adaptation, and genetic conflict in fission yeasts.

## AUTHOR CONTRIBUTIONS

Conceptualization: Guo-Song Jia and Li-Lin Du; Methodology and investigation: Guo-Song Jia, Fang Suo, Ambre Noly, and Philippe Fort; Protein-coding gene manual curation: Guo-Song Jia, Fang Suo, Yue Liang, Wen Li, Wen-Cai Zhang, Heng-Le Li, Xiao-Min Du, Fan-Yi Zhang, and Tong-Yang Du; Writing – review and editing: Guo-Song Jia, Fang Suo, Yu Hua, Feng-Yan Bai, Qi-Ming Wang, Michael Brysch-Herzberg, Dominique Helmlinger, and Li-Lin Du; Funding acquisition: Dominique Helmlinger and Li-Lin Du.

## ACKNOWLEDGEMENTS

This work was supported by the National Key R&D Program of China (2024YFA0917400), the National Natural Science Foundation of China (Grant No. 32470745), and other grants from the Ministry of Science and Technology of China, the Beijing municipal government, and Tsinghua University to L.L.D., and the CNRS MITI 80Prime program to D.H. We thank Drs. Nick Rhind and Huiqiang Lou for their suggestions.

## DATA AVAILABILITY STATEMENT

The data that support the findings of this study are available from the corresponding author upon reasonable request.

---

**Figure S1.**
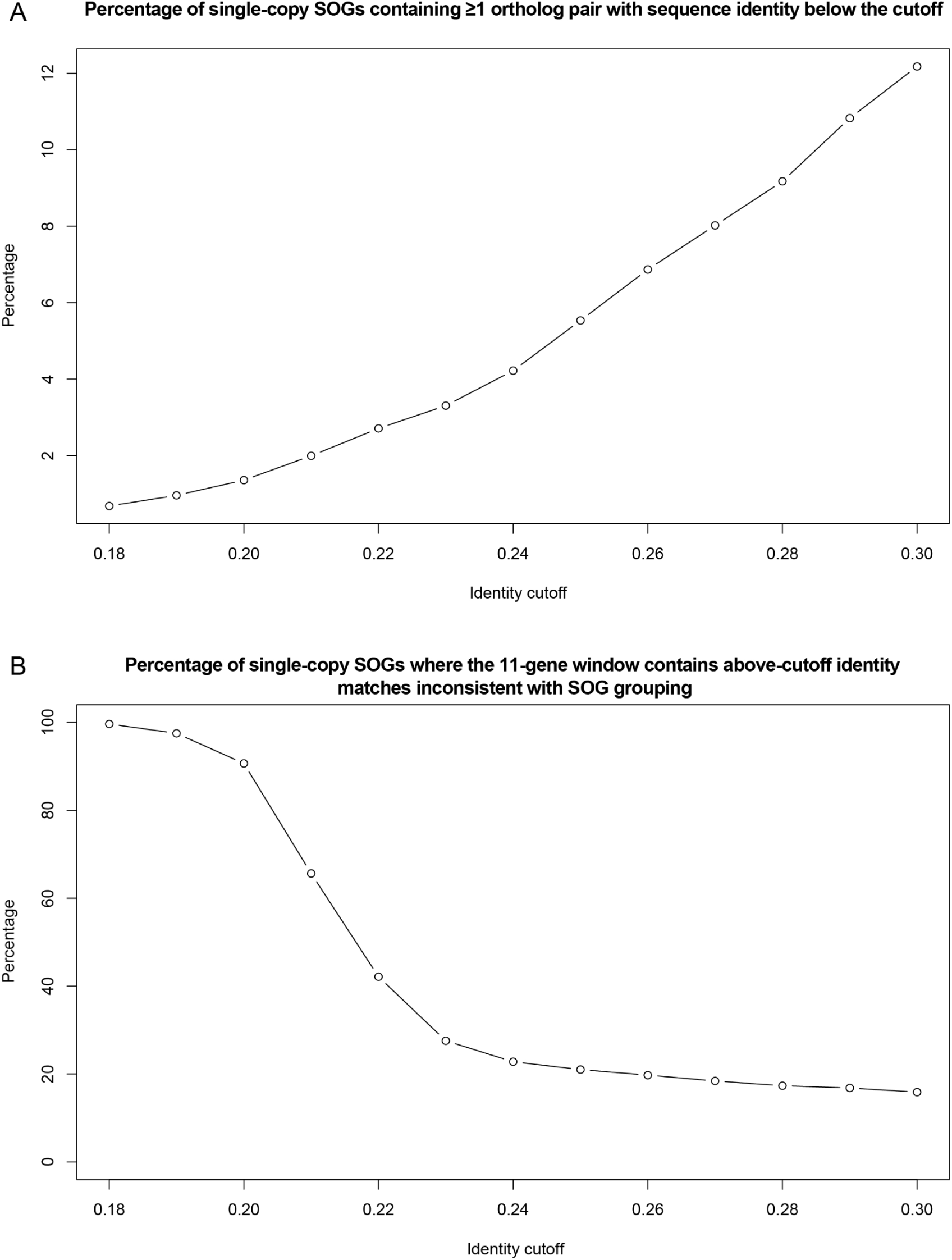
Evaluation of sequence identity thresholds for defining similarity groups. We tested a range of amino acid identity thresholds (0.18 to 0.30, in 0.01 increments) to determine the optimal cutoff for assigning genes to similarity groups within the synteny visualization module. (A) The percentage of single-copy SOGs containing ≥1 ortholog pair with sequence identity below the cutoff. In these instances, true orthologs would fail to be assigned to the same similarity group (false negatives). (B) The percentage of single-copy SOGs where the 11-gene window (the SOG member plus five upstream and five downstream genes) exhibits above-cutoff identity matches inconsistent with SOG assignments. Such matches result in spurious links between non-orthologous genes (false positives).

## REFERENCES

Rhind, N., Chen, Z., Yassour, M., Thompson, D. A., Haas, B. J., Habib, N., … Nusbaum, C. (2011). Comparative functional genomics of the fission yeasts. Science, 332(6032), 930–936. doi:10.1126/science.1203357

Shen, X. X., Steenwyk, J. L., LaBella, A. L., Opulente, D. A., Zhou, X., Kominek, J., … Rokas, A. (2020). Genome-scale phylogeny and contrasting modes of genome evolution in the fungal phylum Ascomycota. Sci Adv, 6(45). doi:10.1126/sciadv.abd0079

Helston, R. M., Box, J. A., Tang, W., & Baumann, P. (2010). Schizosaccharomyces cryophilus sp. nov., a new species of fission yeast. FEMS Yeast Res, 10(6), 779–786. doi:10.1111/j.1567-1364.2010.00657.x

Vaughan-Martini, A., & Martini, A. (2011). Schizosaccharomyces Lindner (1893). In The yeasts (pp. 779–784): Elsevier.

Brysch-Herzberg, M., Tobias, A., Seidel, M., Wittmann, R., Wohlmann, E., Fischer, R., … Peter, G. (2019). Schizosaccharomyces osmophilus sp. nov., an osmophilic fission yeast occurring in bee bread of different solitary bee species. FEMS Yeast Res, 19(4). doi:10.1093/femsyr/foz038

Brysch-Herzberg, M., Jia, G. S., Sipiczki, M., Seidel, M., Li, W., Assali, I., & Du, L. L. (2023). Schizosaccharomyces lindneri sp. nov., a fission yeast occurring in honey. Yeast, 40(7), 237–253. doi:10.1002/yea.3857

Brysch-Herzberg, M., Jia, G. S., Sipiczki, M., Seidel, M., Zhang, W. C., & Du, L. L. (2024). Reinstatement of the fission yeast species Schizosaccharomyces versatilis Wickerham et Duprat, a sibling species of Schizosaccharomyces japonicus. Yeast, 41(3), 108–127. doi:10.1002/yea.3922

Hoffman, C. S., Wood, V., & Fantes, P. A. (2015). An Ancient Yeast for Young Geneticists: A Primer on the Schizosaccharomyces pombe Model System. Genetics, 201(2), 403–423. doi:10.1534/genetics.115.181503

Hayles, J., & Nurse, P. (2018). Introduction to Fission Yeast as a Model System. Cold Spring Harb Protoc, 2018(5). doi:10.1101/pdb.top079749

He, W., Yang, J., Jing, Y., Xu, L., Yu, K., & Fang, X. (2023). NGenomeSyn: an easy-to-use and flexible tool for publication-ready visualization of syntenic relationships across multiple genomes. Bioinformatics, 39(3). doi:10.1093/bioinformatics/btad121

Klar, A. J. (2013). Schizosaccharomyces japonicus yeast poised to become a favorite experimental organism for eukaryotic research. G3 (Bethesda), 3(10), 1869–1873. doi:10.1534/g3.113.007187

Aoki, K., Furuya, K., & Niki, H. (2017). Schizosaccharomyces japonicus: A Distinct Dimorphic Yeast among the Fission Yeasts. Cold Spring Harb Protoc, 2017(12), pdb.top082651. doi:10.1101/pdb.top082651

Seike, T., Shimoda, C., & Niki, H. (2019). Asymmetric diversification of mating pheromones in fission yeast. PLoS Biol, 17(1), e3000101. doi:10.1371/journal.pbio.3000101

Chapman, E., Taglini, F., & Bayne, E. H. (2022). Separable roles for RNAi in regulation of transposable elements and viability in the fission yeast Schizosaccharomyces japonicus. PLoS Genet, 18(2), e1010100. doi:10.1371/journal.pgen.1010100

De Carvalho, M., Jia, G. S., Nidamangala Srinivasa, A., Billmyre, R. B., Xu, Y. H., Lange, J. J., … Zanders, S. E. (2022). The wtf meiotic driver gene family has unexpectedly persisted for over 100 million years. Elife, 11. doi:10.7554/eLife.81149

Rutherford, K. M., Harris, M. A., Oliferenko, S., & Wood, V. (2022). JaponicusDB: rapid deployment of a model organism database for an emerging model species. Genetics, 220(4). doi:10.1093/genetics/iyab223

Alam, S., Gu, Y., Reichert, P., Bähler, J., & Oliferenko, S. (2023). Optimization of energy production and central carbon metabolism in a non-respiring eukaryote. Curr Biol, 33(11), 2175–2186.e2175. doi:10.1016/j.cub.2023.04.046

Gu, Y., Alam, S., & Oliferenko, S. (2023). Peroxisomal compartmentalization of amino acid biosynthesis reactions imposes an upper limit on compartment size. Nat Commun, 14(1), 5544. doi:10.1038/s41467-023-41347-x

Rao, B. D., Gomez-Gil, E., Peter, M., Balogh, G., Nunes, V., MacRae, J. I., … Oliferenko, S. (2025). Horizontal acquisition of prokaryotic hopanoid biosynthesis reorganizes membrane physiology driving lifestyle innovation in a eukaryote. Nat Commun, 16(1), 3291. doi:10.1038/s41467-025-58515-w

Brysch-Herzberg, M., Jia, G. S., Seidel, M., Assali, I., & Du, L. L. (2022). Insights into the ecology of Schizosaccharomyces species in natural and artificial habitats. Antonie Van Leeuwenhoek, 115(5), 661–695. doi:10.1007/s10482-022-01720-0

Jia, G. S., Zhang, W. C., Liang, Y., Liu, X. H., Rhind, N., Pidoux, A., … Du, L. L. (2023). A high-quality reference genome for the fission yeast Schizosaccharomyces osmophilus. G3 (Bethesda), 13(4). doi:10.1093/g3journal/jkad028

Rutherford, K. M., Lera-Ramírez, M., & Wood, V. (2024). PomBase: a Global Core Biodata Resource-growth, collaboration, and sustainability. Genetics, 227(1). doi:10.1093/genetics/iyae007

Lechner, M., Findeiss, S., Steiner, L., Marz, M., Stadler, P. F., & Prohaska, S. J. (2011). Proteinortho: detection of (co-)orthologs in large-scale analysis. BMC Bioinformatics, 12, 124. doi:10.1186/1471-2105-12-124

Emms, D. M., & Kelly, S. (2019). OrthoFinder: phylogenetic orthology inference for comparative genomics. Genome Biol, 20(1), 238. doi:10.1186/s13059-019-1832-y

Lechner, M., Hernandez-Rosales, M., Doerr, D., Wieseke, N., Thévenin, A., Stoye, J., … Stadler, P. F. (2014). Orthology detection combining clustering and synteny for very large datasets. PLoS One, 9(8), e105015. doi:10.1371/journal.pone.0105015

Mullis, A., Lu, Z., Zhan, Y., Wang, T. Y., Rodriguez, J., Rajeh, A., … Lin, Z. (2020). Parallel Concerted Evolution of Ribosomal Protein Genes in Fungi and Its Adaptive Significance. Mol Biol Evol, 37(2), 455–468. doi:10.1093/molbev/msz229

Katoh, K., & Standley, D. M. (2013). MAFFT multiple sequence alignment software version 7: improvements in performance and usability. Mol Biol Evol, 30(4), 772–780. doi:10.1093/molbev/mst010

Minh, B. Q., Schmidt, H. A., Chernomor, O., Schrempf, D., Woodhams, M. D., von Haeseler, A., & Lanfear, R. (2020). IQ-TREE 2: New Models and Efficient Methods for Phylogenetic Inference in the Genomic Era. Mol Biol Evol, 37(5), 1530–1534. doi:10.1093/molbev/msaa015

Paradis, E., & Schliep, K. (2019). ape 5.0: an environment for modern phylogenetics and evolutionary analyses in R. Bioinformatics, 35(3), 526–528. doi:10.1093/bioinformatics/bty633

Gilchrist, C. L. M., & Chooi, Y. H. (2021). clinker & clustermap.js: automatic generation of gene cluster comparison figures. Bioinformatics, 37(16), 2473–2475. doi:10.1093/bioinformatics/btab007

Butler, M., Goodwin, T., Simpson, M., Singh, M., & Poulter, R. (2001). Vertebrate LTR retrotransposons of the Tf1/sushi group. J Mol Evol, 52(3), 260–274. doi:10.1007/s002390010154

Goodwin, T. J., & Poulter, R. T. (2001). The diversity of retrotransposons in the yeast Cryptococcus neoformans. Yeast, 18(9), 865–880. doi:10.1002/yea.733

Lin, Z., & Li, W. H. (2011). Expansion of hexose transporter genes was associated with the evolution of aerobic fermentation in yeasts. Mol Biol Evol, 28(1), 131–142. doi:10.1093/molbev/msq184

Karavirta, V., & Shaffer, C. A. (2013). JSAV: the JavaScript algorithm visualization library. Paper presented at the Annual Conference on Innovation and Technology in Computer Science Education.

Shank, S. D., Weaver, S., & Kosakovsky Pond, S. L. (2018). phylotree.js - a JavaScript library for application development and interactive data visualization in phylogenetics. BMC Bioinformatics, 19(1), 276. doi:10.1186/s12859-018-2283-2

Taricani, L., Feilotter, H. E., Weaver, C., & Young, P. G. (2001). Expression of hsp16 in response to nucleotide depletion is regulated via the spc1 MAPK pathway in Schizosaccharomyces pombe. Nucleic Acids Res, 29(14), 3030–3040. doi:10.1093/nar/29.14.3030

Hirose, M., Tohda, H., Giga-Hama, Y., Tsushima, R., Zako, T., Iizuka, R., … Yohda, M. (2005). Interaction of a small heat shock protein of the fission yeast, Schizosaccharomyces pombe, with a denatured protein at elevated temperature. J Biol Chem, 280(38), 32586–32593. doi:10.1074/jbc.M504121200

Yoshida, J., & Tani, T. (2005). Hsp16p is required for thermotolerance in nuclear mRNA export in fission yeast Schizosaccharomyces pombe. Cell Struct Funct, 29(5-6), 125–138. doi:10.1247/csf.29.125

Glatz, A., Pilbat, A. M., Németh, G. L., Vince-Kontár, K., Jósvay, K., Hunya, Á., … Török, Z. (2016). Involvement of small heat shock proteins, trehalose, and lipids in the thermal stress management in Schizosaccharomyces pombe. Cell Stress Chaperones, 21(2), 327–338. doi:10.1007/s12192-015-0662-4

Tong, P., Pidoux, A. L., Toda, N. R. T., Ard, R., Berger, H., Shukla, M., … Allshire, R. C. (2019). Interspecies conservation of organisation and function between nonhomologous regional centromeres. Nat Commun, 10(1), 2343. doi:10.1038/s41467-019-09824-4

Haslbeck, M., & Vierling, E. (2015). A first line of stress defense: small heat shock proteins and their function in protein homeostasis. J Mol Biol, 427(7), 1537–1548. doi:10.1016/j.jmb.2015.02.002

Waters, E. R., & Vierling, E. (2020). Plant small heat shock proteins - evolutionary and functional diversity. New Phytol, 227(1), 24–37. doi:10.1111/nph.16536

Xu, Y. H., Suo, F., Zhang, X. R., Du, T. Y., Hua, Y., Jia, G. S., … Du, L. L. (2024). Evolutionary Modes of wtf Meiotic Driver Genes in Schizosaccharomyces pombe. Genome Biol Evol, 16(10). doi:10.1093/gbe/evae221

Zhang, F.-Y., Jia, G.-S., Ren, J.-Y., Suo, F., Du, T.-Y., Zhang, W.-C., … Hua, Y. (2025). Evolutionary persistence and divergence of the tdk killer meiotic driver family. 2025.2012.2028.696746. doi:10.64898/2025.12.28.696746 bioRxiv

